# Evaluation of *de novo* transcriptome assemblies from RNA-Seq data

**DOI:** 10.1101/006338

**Authors:** Bo Li, Nathanael Fillmore, Yongsheng Bai, Mike Collins, James A. Thomson, Ron Stewart, Colin N. Dewey

## Abstract

*De novo* RNA-Seq assembly facilitates the study of transcriptomes for species without sequenced genomes, but it is challenging to select the most accurate assembly in this context. To address this challenge, we developed a model-based score, RSEM-EVAL, for evaluating assemblies when the ground truth is unknown. Our experiments show that RSEM-EVAL correctly reflects assembly accuracy, as measured by REF-EVAL, a refined set of ground-truth-based scores that we also developed. With the guidance of RSEM-EVAL, we assembled the transcriptome of the regenerating axolotl limb; this assembly compares favorably to a previous assembly.

## Background

RNA-Seq technology is revolutionizing the study of species that have not yet had their genomes sequenced by enabling the large-scale analysis of their transcriptomes. In order to study such transcriptomes, one must first determine a set of transcript sequences via *de novo* transcriptome assembly, the task of reconstructing transcript sequences from RNA-Seq reads without the aid of genome sequence information. A number of *de novo* transcriptome assemblers are currently available, many designed for Illumina platform data [2, 4, 12, 27, 35, 37, 40, 45] and others targeted for Roche 454 Life Science platform data [3, 15, 26, 49]. These assemblers, combined with their often sizable sets of user-tunable parameters, enable the generation of a large space of candidate assemblies for a single data set. However, appropriately evaluating the accuracy of assemblies in this space, particularly when the ground truth is unknown, has remained challenging.

A number of recent studies have been devoted to the evaluation of *de novo* transcriptome assemblies [6, 10, 18, 24, 29, 31, 34, 48]. Assembly evaluation measures used in such studies can be grouped into two classes: reference-based and reference-free. Reference-based measures are those that are computed using previously known sequences. For example, after establishing a correspondence between assembly elements and a reference transcript set, one can calculate the fraction of assembly elements that accurately match a reference transcript (precision), the fraction of reference transcripts that are matched by assembly elements (recall), or a combination of these two (e.g., the *F*_1_-measure) [2, 29, 34]. In addition to transcript sets, genome and protein sequences can also be used as references for assembly evaluation [12, 18, 35, 45, 48].

However, in most cases where *de novo* assembly is of interest, reference sequences are either not available, incomplete, or considerably diverged from the ground truth of a sample of interest, which makes the assembly evaluation task markedly more difficult. In such cases, one must resort to reference-free measures. Commonly used reference-free measures include median contig length, number of contigs, and N50 [18, 29, 34]. Unfortunately, these measures are primitive and often misleading. For example, N50, one of the most popular reference-free measures, can be maximized by trivial assemblies. N50 is defined as the length of the longest contig such that all contigs of at least that length compose at least 50% of the bases of the assembly [28]. The motivation for this measure is that better assemblies will result from a larger number of identified overlaps between the input reads and thus will have more reads assembled into longer contigs. However, it is easy to see that a trivial assembly constructed by concatenating all of the input reads into a single contig will maximize this measure. In short, N50 measures the continuity of contigs but not their accuracy [36]. Other simplistic reference-free measures can be similarly misleading regarding the accuracy of assemblies.

We improve upon the state-of-the-art in transcriptome assembly evaluation by presenting the DETONATE (*DE novo* TranscriptOme rNa-seq Assembly with or without the Truth Evaluation) methodology and software package. DETONATE consists of two components: RSEM-EVAL and REF-EVAL. RSEM-EVAL, DETONATE’s primary contribution, is a reference-free evaluation method based on a novel probabilistic model that depends only on an assembly and the RNA-Seq reads used to construct it. RSEM-EVAL is similar to recent approaches using statistical models to evaluate or construct genome [33] and metagenome [5, 20] assemblies, but, as we will discuss, is necessarily more complex because of widely-varying abundances of transcripts and alternative splicing. Unlike N50, RSEM-EVAL combines multiple factors, including the compactness of an assembly and the support of the assembly from the RNA-Seq data, into a single, statistically-principled evaluation score. This score can be used to select a best assembler, optimize an assembler’s parameters, and guide new assembler design as an objective function. In addition, for each contig within an assembly, RSEM-EVAL provides a score that assesses how well that contig is supported by the RNA-Seq data and can be used to filter unnecessary contigs. REF-EVAL, DETONATE’s second component, is a toolkit of reference-based measures. It provides a more refined view of an assembly’s accuracy than existing reference-based measures.

We have performed a number of experiments on both real and simulated data to demonstrate the value of the RSEM-EVAL score. First, we generated a set of perturbed assemblies around a single “true” assembly, and we show that RSEM-EVAL ranks the truth among the highest scoring assemblies. Second, we computed RSEM-EVAL scores and REF-EVAL reference-based measures on over 200 assemblies for multiple data sets, and we find that the RSEM-EVAL score generally correlates well with reference-based measures. The results of these first two experiments together show that the RSEM-EVAL score accurately evaluates *de novo* transcriptome assemblies, despite not having knowledge of the ground truth. Lastly, as a demonstration of the use of the RSEMEVAL score, we assembled the transcriptome of the regenerating axolotl limb with its guidance. This new assembly allowed for the identification of many more genes that are involved in the process of axolotl limb regeneration than had been found with an assembly from a previous study.

## Results

### DETONATE: a software package for the evaluation of *de novo* transcriptome assemblies

The main contribution of our work is DETONATE, a methodology for the evaluation of *de novo* transcriptome assemblies and a software package that realizes this methodology. DETONATE consists of two components: RSEM-EVAL, which does not require a ground truth reference, and REF-EVAL, which does. The high-level workflow of DETONATE is shown in Figure 1. In the following subsections, we (1) describe the RSEM-EVAL reference-free score and show that the “true” assembly is an approximate local maximum of this score, (2) describe the REF-EVAL reference-based scores and show that RSEM-EVAL’s score correlates well with these measures, indicating that RSEM-EVAL reflects the accuracy of a transcriptome assembly, and (3) demonstrate our methods’ practical usefulness by assembling the transcriptome of the regenerating axolotl limb with the guidance of RSEM-EVAL. To best understand the components of DETONATE and the experiments we have performed with them, we first define what is considered to be the “true” assembly of a set of RNA-Seq reads, which is critical to the assembly evaluation task.

**Figure 1:**
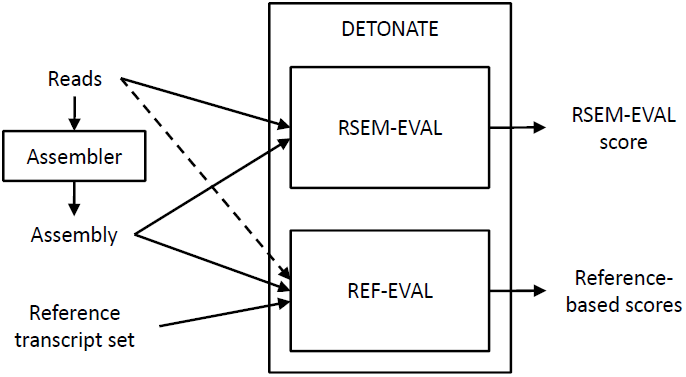
The DETONATE package workflow. The DETONATE package consists of two components: RSEM-EVAL and REF-EVAL. Combined, these two components allow for the computation of a variety of evaluation scores for a *de novo* transcriptome assembly. RSEM-EVAL produces an evaluation score that is based only on an assembly and the set of reads from which it was constructed. When a reference transcript set is available, REF-EVAL may be used to compute a number of reference-based measures. For most measures, REF-EVAL requires only an assembly and a reference transcript set. For weighted measures and measures with respect to an estimated “true” contig set, REF-EVAL additionally requires the set of reads that were assembled (dashed arrow).

### The “true” assembly according to DETONATE

Ideally, the goal of transcriptome assembly would be to construct the full-length sequences of all transcripts in the transcriptome. Unfortunately, it is rarely possible to acheive this goal in practice, because sequencing depths are usually not high enough to completely cover all transcripts, especially those of low abundance. Thus, a transcriptome assembly is, in general, a set of *contigs*, with each contig representing the sequence of a segment of a transcript. When paired-end data are assembled, one can also consider constructing *scaffolds*, or chains of contigs separated by unknown sequences with estimated lengths. In this work, we restrict our attention to contig assemblies constructed from single-end data.

Both RSEM-EVAL and REF-EVAL make use of the concept of the “true” assembly of a set of RNA-Seq reads, which is the assembly one would construct if given knowledge of the true origin of each read. A precise definition of the “true” assembly is provided in the Methods. Intuitively, for each nonnegative integer *w*, the “true” assembly at minimum overlap length *w* is the collection of transcript subsequences that are covered by reads whose true alignments overlap by at least *w* bases. The “true” assembly at minimum overlap length 0 is the collection of transcript subsequences covered by reads whose true alignments are contiguous (that is, overlap by 0 bases) or overlap by at least one base. The “true” assembly at minimum overlap length 0 is the best theoretically achievable assembly in that it represents all supported segments of the transcript sequences, and no unsupported segments. Figure 2 gives an example of “true” assemblies at minimum overlap lengths *w* = 0 and *w* = 1.

**Figure 2:**
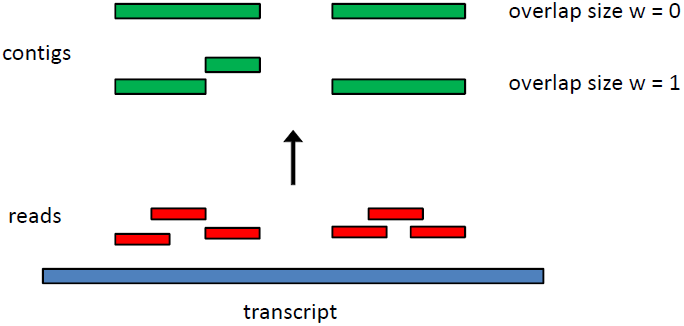
An example construction of “true” assemblies with different minimum overlap lengths. Six reads (red) are positioned at their true places of origin along one transcript (blue). The assembly with minimum overlap length *w* = 0 consists of two contigs (green). For *w* = 1, the assembly instead consists of three contigs because the second and third reads from the left are immediately adjacent but not overlapping.

### RSEM-EVAL is a novel reference-free transcriptome assembly evaluation measure

RSEM-EVAL, our primary contribution, is a reference-free evaluation measure based on a novel probabilistic model that depends only on an assembly and the RNA-Seq reads used to construct it. In short, the RSEM-EVAL score of an assembly is defined as the log joint probability of the assembly *A* and the reads *D* used to construct it, under a model that we have devised. In symbols,

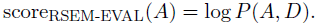

RSEM-EVAL’s intended use is to compare several assemblies of the same set of reads, and under this scenario, the joint probability is proportional to the posterior probability of the assembly given the reads. Although the posterior probability of the assembly given the reads is a more natural measure for this application, we use the joint probability because it is more efficient to compute.

The details of the RSEM-EVAL model are provided in the Methods. In summary, the RSEM-EVAL score consists of three components: an assembly prior, a likelihood, and a BIC penalty. That is,

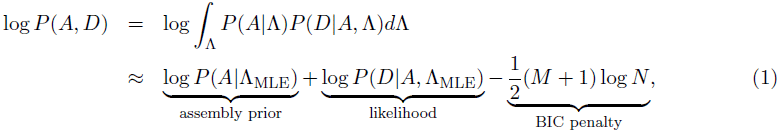

where *N* is the total number of reads, *M* is the number of contigs in the assembly, and Λ_MLE_ is the maximum likelihood (ML) estimate of the expected read coverage under *P* (*A, D|*Λ). For typical sizes of RNA-Seq data sets used for transcriptome assembly, the likelihood is generally the dominant component of the RSEM-EVAL score in the above equation. It serves to assess how well the assembly explains the RNA-Seq reads. However, as we will show later, only having this component is not enough. Thus we use the assembly prior and BIC components to assess an assembly’s complexity. These two components penalize assemblies with too many bases or contigs, or with an unusual distribution of contig lengths relative to the expected read coverage. Thus, these two components impose a parsimony preference on the RSEM-EVAL score. The three components together enable the RSEM-EVAL score to favor simple assemblies that can explain the RNA-Seq reads well.

### The ground truth is an approximate local maximum of the RSEM-EVAL score

As we have discussed, the “true” assembly at minimum overlap length 0 is the best possible assembly that can be constructed solely from RNA-Seq data. Therefore, we consider it to be the ground truth assembly. A good evaluation measure should score the ground truth among its best assemblies. Ideally, we would have explored the entire space of assemblies and shown that the RSEM-EVAL score for the ground truth assembly is among the highest scores. However, such a global search of assembly space is computationally infeasible. Thus, we instead performed experiments that assess whether in the local space around the ground truth, the ground truth is among the best scoring assemblies. In other words, we tested whether the ground truth assembly is approximately a *local maximum* of the RSEM-EVAL function.

We explored the local space of assemblies around that of the ground truth by generating assemblies that were slightly perturbed from it. We performed experiments with two kinds of perturbations: random perturbations and guided perturbations. In our random perturbation experiments, assemblies were generated by randomly mutating the ground truth assembly. Since the minimum overlap length is a critical parameter for constructing assemblies, we also assessed the RSEM-EVAL scores for “true” assemblies with different minimum overlap lengths in guided perturbation experiments. A good evaluation score should generally prefer “true” assemblies with small minimum overlap lengths, which are closest to the ground truth.

For these experiments, it was critical that the ground truth assembly be known, and therefore we primarily used a simulated set of RNA-Seq data, in which the true origin of each read is known. In addition, for our guided perturbation experiments, we used the real mouse data set on which the simulated data were based, and we estimated the true origin of each read. For details about these real and simulated data, see the Methods.

### Random perturbation

With our random perturbation experiment we wished to determine how well, in terms of the RSEMEVAL score, the ground truth compares to assemblies in the local space surrounding it. In order to thoroughly explore the space of assemblies centered at the ground truth, we used four types of mutations (substitution, fusion, fission, and indels), each of which was applied at five different strength levels (see Methods). Therefore, in total, we generated 20 classes of randomly perturbed assemblies. For each class, we generated 1000 independent randomly perturbed assemblies in order to estimate the RSEM-EVAL score population mean and its 95% confidence interval for assemblies of that class.

On average, the perturbed assemblies had RSEM-EVAL scores that were worse than that of the ground truth (Figure 3(A)). In addition, the higher the mutation strength, the worse the mean score of the perturbed assemblies. This suggests that the ground truth assembly behaves similarly to a local maximum of the RSEM-EVAL function. Even though the population mean scores of the perturbed assemblies were estimated to be always worse than the score of the ground truth, individual perturbed assemblies could have had higher scores. Therefore, for each class of perturbed assemblies, we calculated the fraction of assemblies with RSEM-EVAL scores larger than that of the ground truth, which we refer to as the *error rate*. Error rates decreased dramatically with increasing mutation strength and, for all mutation types except fusion, error rates were only non-zero for the weakest mutation strength level (Figure 3(B)). RSEM-EVAL had the most trouble with the fusion-perturbed assemblies, with more than half of such assemblies at the weakest mutation strength having a score above that of the ground truth. From individual examinations of these assemblies, we observed that in many of these cases, the assemblies contained fusions of contigs with low abundances, which are difficult to distinguish from true contigs, especially with the ground truth defined as the “true” assembly with minimum overlap length *w* = 0.

**Figure 3:**
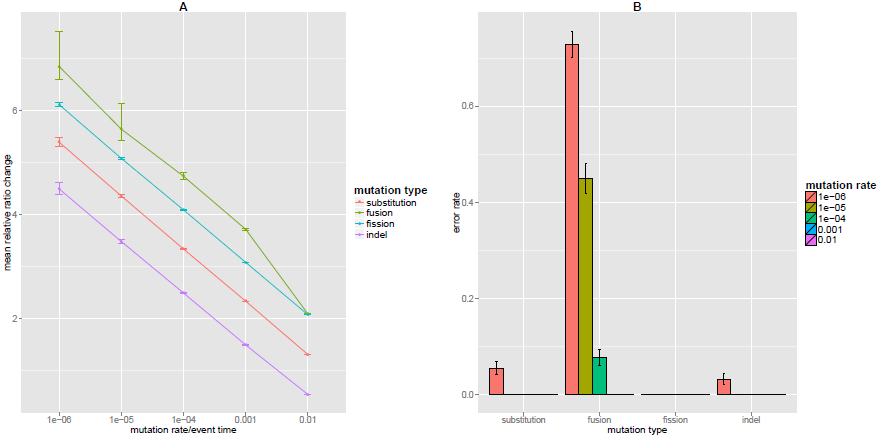
Random perturbation results. Comparison of the RSEM-EVAL score of the ground truth assembly to those of randomly perturbed versions of that assembly. (A) Changes in the relative scores of the perturbed assemblies with increasing mutation strength. For each class of perturbed assemblies, we computed the mean relative ratio change, *ρ*, which is defined as 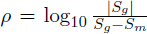, where *S_g_* is the RSEM-EVAL score for the ground truth assembly and *S_m_* is the mean RSEM-EVAL score for the 1000 randomly-perturbed assemblies in that class. Smaller (worse) mean scores correspond to smaller values for *ρ*. For each mutation type, *ρ* is plotted as a function of the mutation strength, with error bars corresponding to 95% confidence intervals for *ρ*. (B) RSEM-EVAL error rates for each perturbed assembly class. Error bars correspond to the 95% confidence intervals for the mean error rates.

### Guided perturbation

With the guided perturbation experiments, we measured the RSEM-EVAL scores of assemblies constructed with different values of the minimum overlap length, which is a common parameter in assemblers. Since the “true” assembly at minimum overlap length 0 is the best achievable assembly, a good evaluation score should be maximized at small minimum overlap lengths. As described before, we used one simulated and one real mouse RNA-Seq data set for these experiments. For each data set, we constructed the “true” assemblies with minimum overlap lengths ranging from 0 to 75. The “true” assembly at minimum overlap length 76 (the read length) was not constructed because of prohibitive running times. For the real RNA-Seq data, “true” assemblies were estimated using REF-EVAL’s procedure, described below. We then calculated the RSEM-EVAL scores for all of these assemblies.

As we had hoped, we found that the RSEM-EVAL score was maximized at small minimum overlap lengths, for both the simulated and real data sets (Figure 4). In contrast, the ML score increased with minimum overlap length and peaked at minimum overlap length *w* = 75. These results support the necessity of the prior component of the RSEM-EVAL model, which takes into account the complexity of an assembly.

**Figure 4:**
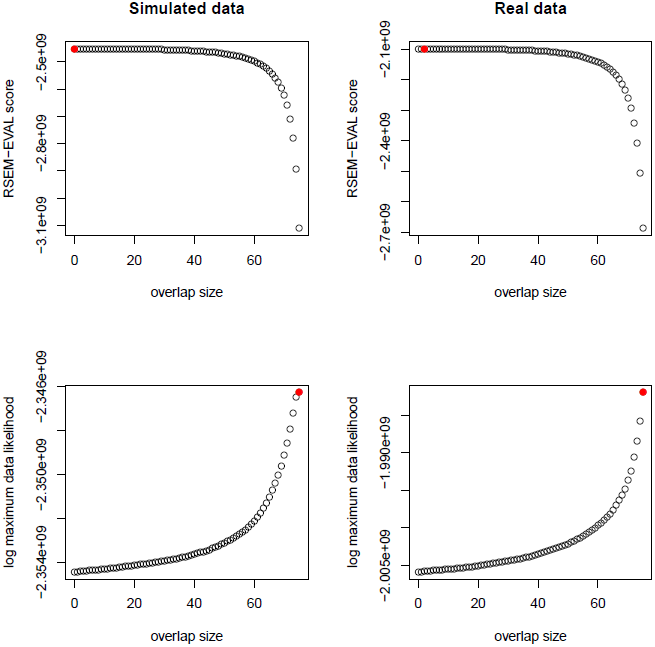
Guided perturbation results. RSEM-EVAL (top row) and ML (bottom row) scores of “true” assemblies for different values of the minimum overlap length *w* on both simulated (left column) and real (right column) data sets. The maximizing values (red circles) are achieved at *w* = 0, *w* = 2, *w* = 75 and *w* = 75 in a top-down, left-right order. For better visualization of the maximizing values of *w*, RSEM-EVAL scores for the local regions around the maximal values are shown in Additional file 1, Figure S2.

To explore the effects of the minimum overlap length parameter, *w*, in the RSEM-EVAL model, we also performed the guided perturbation experiments with *w* = 50 for the RSEM-EVAL model. We did not observe any major differences between these results and those for *w* = 0 (Additional file 1, Figure S3, S4). In order to explain this result, note that although the RSEM-EVAL model uses *w* in both the prior and likelihood correction components, our estimation procedure for the uncorrected likelihood component (see Methods) does not explicitly check for violations of the minimum overlap length by the assembly (i.e., regions that are not covered by reads that overlap each other by at least *w* bases). Thus, the minimum overlap length does not play a role in the uncorrected likelihood, which is the dominant term of the RSEM-EVAL score.

### REF-EVAL is a refined toolset for computing reference-based evaluation measures

Our first experiment, above, shows that RSEM-EVAL has an approximate local maximum at the “true” assembly. However, this does not necessarily imply that RSEM-EVAL induces a useful ranking of assemblies away from this local maximum.

Thus, in order to assess the usefulness of RSEM-EVAL’s reference-free score, it is of interest to compare the ranking RSEM-EVAL assigns to a collection of assemblies to the ranking assigned by comparing each assembly to a reference. This raises two questions: (1) what reference to compare against, and (2) how to perform the comparison. REF-EVAL constitutes an answer to both questions. The tools REF-EVAL provides are also of independent interest for reference-based evaluation of transcriptome assemblies.

In answer to question (1), REF-EVAL provides a method to estimate the “true” assembly of a set of reads, relative to a collection of full-length reference transcript sequences. The estimate is based on alignments of reads to reference transcripts, as described in the Methods. As we have previously discussed, we wish to compare assemblies against the set of “true” contigs instead of full-length reference sequences because the latter cannot, in general, be fully reconstructed from the data and we want to reward assemblies for recovering read-supported subsequences of the references.

In answer to question (2), REF-EVAL provides two kinds of reference-based measures. First, REFEVAL provides assembly recall, precision, and *F*_1_ scores at two different granularities: contig and nucleotide. Recall is the fraction of reference elements (contigs or nucleotides) that are correctly recovered by an assembly. Precision is the fraction of assembly elements that correctly recover a reference element. The *F*_1_ score is the harmonic mean of recall and precision. For precise definitions and computational details, see Methods.

Although the contig- and nucleotide-level measures are straightforward and intuitive, both have drawbacks and the two measures can be quite dissimilar (Figure 5). For example, if two contigs align perfectly to a single reference sequence, but neither covers at least 99% of that sequence, the nucleotide-level measure will count them as correct, whereas the contig-level measure will not (Figure 5(B)). In general, the contig-level measurement can fail to give a fair assessment of an assembly’s overall quality, since it uses very stringent criteria and normally only a small fraction of the reference sequences are correctly recovered. And whereas the nucleotide-level measure arguably gives a more detailed picture of an assembly’s quality, it fails to take into account connectivity between nucleotides. For example, in the example depicted in Figure 5(B), the nucleotide-level measure does not take into account the fact that the correctly predicted nucleotides of the reference sequence are predicted by two different contigs.

**Figure 5:**
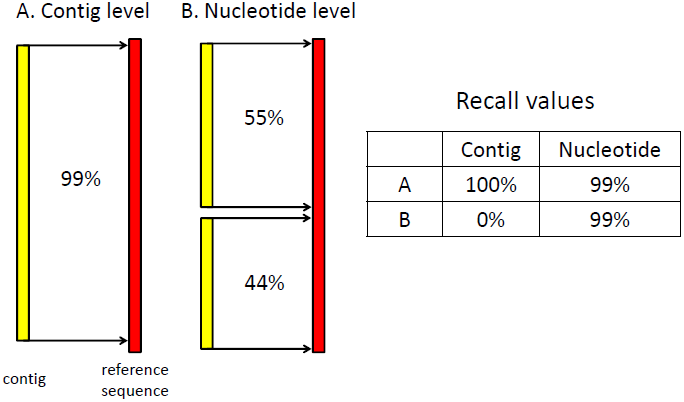
The different granularities of reference-based measures computed by REF-EVAL. (A) The contig level measure requires at least 99% alignment between a matched contig and reference sequence in a one-to-one mapping between an assembly and the reference. (B) The nucleotide level measure counts the number of correctly recovered nucleotides without requiring a one-to-one mapping. Unlike the contig level measure, it gives full credit to the two short contigs. The table on the right gives both the contig-level and nucleotide-level recall values for (A) and (B).

Motivated, in part, by the shortcomings of the contig- and nucleotide-level measures, REF-EVAL also provides a novel transcriptome assembly reference-based accuracy measure, the *k*-mer compression score (KC score). In devising the KC score, our goals were to define a measure that would (1) address some of the limitations of the contig- and nucleotide-level measures, (2) provide further intuition for what the RSEM-EVAL score optimizes, and (3) be relatively simple. The KC score is a combination of two measures, weighted *k*-mer recall (WKR) and inverse compression rate (ICR), and is simply defined as

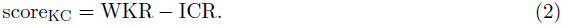

The WKR measures the *fidelity* with which a particular assembly represents the *k*-mer content of the reference sequences. Balancing the WKR, the ICR measures the degree to which the assembly *compresses* the RNA-Seq data. The WKR and ICR are defined and further motivated in the Methods.

### The RSEM-EVAL score correlates highly with reference-based measures

Having specified a framework for reference-based transcriptome assembly evaluation via REF-EVAL, we then sought to test whether the RSEM-EVAL score ranks assemblies similarly to REF-EVAL’s reference-based measures. To test this, we constructed a large number of assemblies on several RNA-Seq data sets from organisms for which reference transcript sequences were available, and we computed both the RSEM-EVAL score and reference-based measures for each assembly. The RNA-Seq data sets used were the simulated and real mouse strand non-specific data from the perturbation experiments, a real strand-specific mouse data set, and a real strand-specific yeast data set. Four publicly available assemblers, Trinity [12], Oases [37], SOAPdenovo-Trans [45] and Trans-ABySS [35], were applied to assemble these data sets using a wide variety of parameter settings.

### Overall correlation

For each data set, we computed Spearman’s rank correlation between the reference-based measure values and the RSEM-EVAL scores to measure the similarity of the rankings implied by them. RSEM-EVAL scores had decent correlation with the contig and nucleotide-level *F*_1_ measures on both the strand non-specific (Figure 6) and strand-specific (Additional file 1, Figure S5) data sets. Specifically, the correlation of the contig and nucleotide-level *F*_1_ measures to the RSEM-EVAL score is comparable to the correlation of the contig and nucleotide-level *F*_1_ measures to each other. In contrast, N50 had poor correlation with these reference-based measures, particularly at the nucleotide level (Additional file 1, Figure S6, S7).

**Figure 6:**
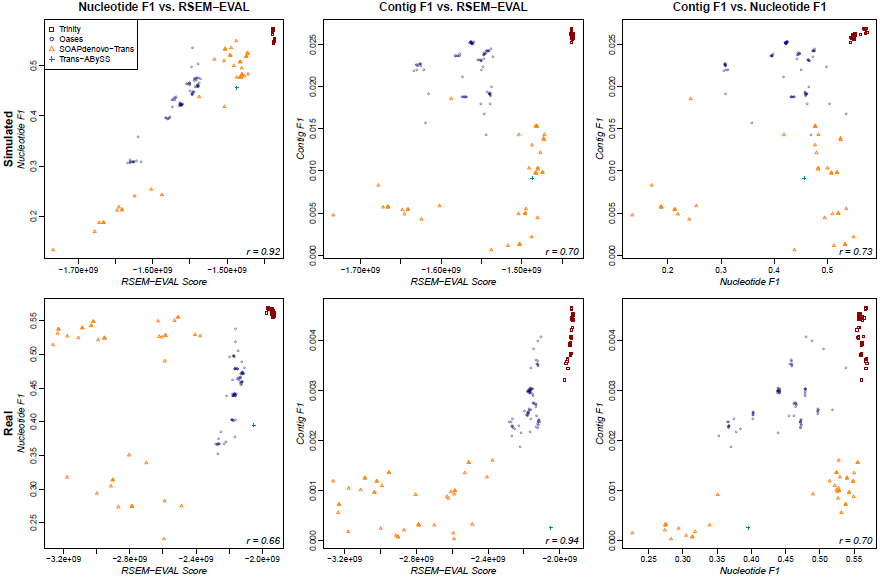
Correlation of the RSEM-EVAL score with reference-based measures on the strand non-specific data sets. Scatterplots are shown for the simulated (top row) and real mouse (bottom row) data sets and for both the nucleotide-level *F*_1_ (left column) and contig-level *F*_1_ (center column) measures. For comparison, scatterplots of the nucleotide-level *F*_1_ against the contig-level *F*_1_ are shown (right column). The Spearman rank correlation coefficient (bottom-right corner of each plot) was computed for each combination of data set and reference-based measure.

The RSEM-EVAL scores had markedly higher correlations with the KC score (*k* = *L*, the read length) for both the strand non-specific (Figure 7) and strand-specific (Additional file 1, Figure S8) data sets, which confirmed our expectations given the mathematical connections between these scores. As with the contig and nucleotide-level measures, N50 similarly had poor correlation with the KC score (Additional file 1, Figure S9, S10). In order to assess the impact of the *k*-mer size on the KC score, we also computed correlations between the RSEM-EVAL score and the KC score at half (*k* = 36) and double (*k* = 152) the read length for the strand non-specific data. We found that these correlation values were not sensitive to the value of *k* (Additional file 1, Figure S11, S12). These results provide some intuition for what the RSEM-EVAL score assesses and indicate that the RSEM-EVAL score could be used as a proxy for the KC score when reference sequences are not known.

**Figure 7:**
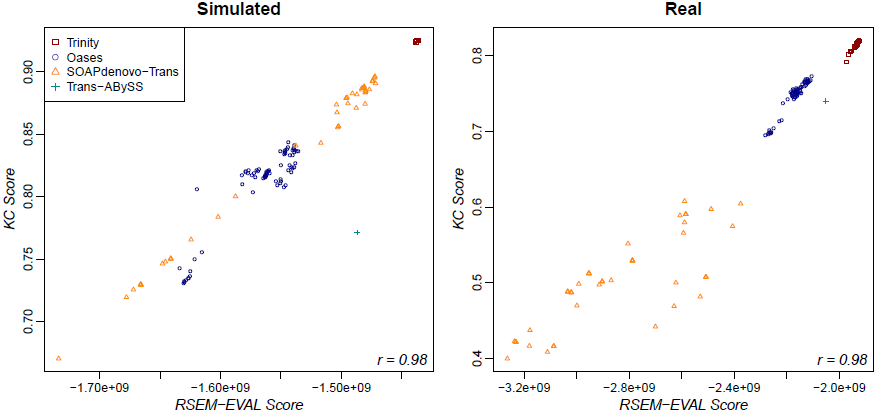
Correlation of the RSEM-EVAL and KC scores on the strand non-specific data sets. The Spearman rank correlation coefficient (bottom-right corner of each plot) was computed for each data set.

Although this experiment was not designed as a comprehensive evaluation, some features of these results are suggestive of the relative accuracies of the assemblers. First, given the selected assembler versions and parameter settings, Trinity produced the most accurate assemblies on all data sets with respect to the contig- and nucleotide-level *F*_1_ scores and the KC score. Unlike N50, the RSEM-EVAL score supported this result, with the Trinity assemblies also obtaining the highest RSEM-EVAL scores. Second, varying the parameters of Trinity had a relatively small effect on the accuracy of the resulting assemblies, as compared to Oases and SOAPdenovo-Trans, which produced assemblies that spanned a large range of accuracies.

### Within-assembler correlation

One important potential application of RSEM-EVAL is the optimization of the parameters of an assembler. Thus, it is of interest whether the RSEM-EVAL score correlates well with reference-based measures for assemblies generated by a single assembler. In the previous subsection, we showed that the RSEM-EVAL score has high correlation with the KC score on real and simulated data when several different assemblers are used. Looking at each assembler separately, we also find that the RSEM-EVAL score has high correlation with the KC score when only the assembler’s parameters are changed, for both strand non-specific (Figure 8) and strand-specific (Additional file 1, Figure S13) data sets. This suggests that RSEM-EVAL can be used to optimize the parameters of an assembler for a given data set when the KC score is of interest for measuring the accuracy of an assembly.

**Figure 8:**
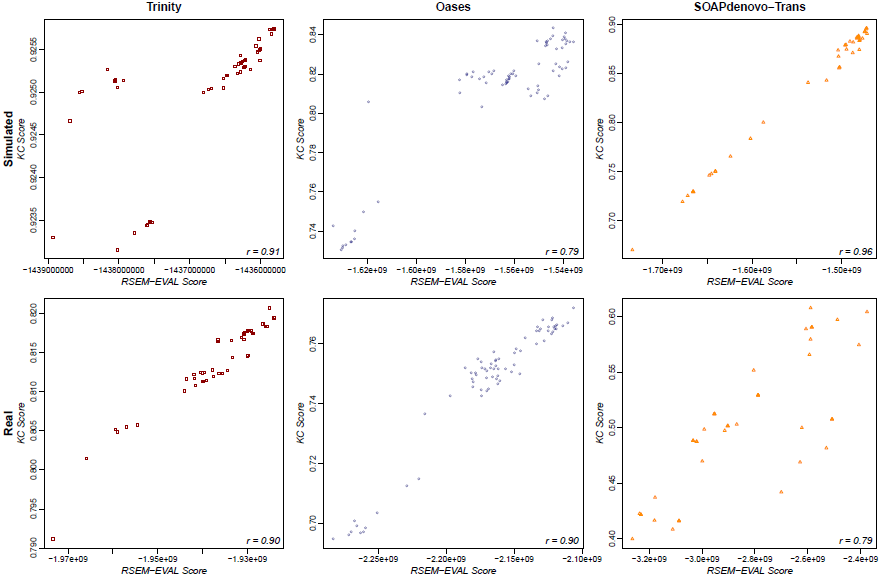
Within-assembler correlation of the RSEM-EVAL and KC scores on the strand non-specific data sets. Scatterplots are shown for the simulated (top row) and real mouse (bottom row) data sets and for the Trinity (left column), Oases (center column), and SOAPdenovo-Trans (right column) assemblers. TransABySS was omitted because it had only one assembly. The Spearman rank correlation coefficient (bottom-right corner of each plot) was computed for each combination of data set and assembler.

### Assessing the relative impact of individual contigs within an assembly

The RSEM-EVAL score is an assembly-level measure that allows one to compare different assemblies constructed from the same data set. It is also of interest to compute scores for individual contigs within an assembly that reflect their relative impacts. One natural way to do this statistically is to compare the hypothesis that a particular contig is a “true” contig with the null hypothesis that the reads composing the contig are actually from the background noise. For each contig, we use the log of the ratio between the probabilities for these two hypotheses as its *contig impact score*. Through a decomposition of the RSEM-EVAL score log *P* (*A, D*) into contig-level components, we are able to calculate these contig impact scores efficiently (Additional file 1, Section 5).

RSEM-EVAL’s contig impact score measures the relative contribution of each contig to explaining the assembled RNA-Seq data. This suggests a strategy to improve the accuracy of an assembly: “trim” it by removing contigs that contribute little to explaining the data. To evaluate this strategy (and by extension the contig impact score itself), we trimmed assemblies of the simulated data using the RSEM-EVAL contig impact scores and computed the resulting changes in the evaluation measures. Assemblies were trimmed by removing all contigs with negative scores.

In general, the trimmed assemblies had better evaluation scores than their untrimmed counterparts (Additional file 1, Table S2 and Figure S14). The largest improvements were seen for assemblies produced by Oases and Trans-ABySS, which tend to produce large numbers of contigs. In fact, for both the nucleotide- and contig-level *F*_1_ scores, the trimmed Oases assemblies were the most accurate of all assemblies (both trimmed and untrimmed), supporting the usefulness of the RSEMEVAL contig impact score. This suggests that the RSEM-EVAL contig impact scores are correctly identifying contigs that are either erroneous or redundant within these assemblies.

### RSEM-EVAL guides creation of an improved axolotl assembly

The axolotl (*Ambystoma mexicanum*) is a neotenic salamander with regenerative abilities that have piqued the interests of scientists. In particular, there is significant interest in studying the molecular basis of axolotl limb regeneration [39]. Although the axolotl is an important model organism, its genome is large and repetitive, and, as a result, it has not yet been sequenced. In addition, a publicly available, complete and high quality set of axolotl transcript sequences does not exist, which makes it challenging to study the axolotl’s transcriptome.

To demonstrate the use of RSEM-EVAL, we employed it to select an assembler and parameter values for a set of previously published RNA-Seq data from a timecourse study of the regenerating axolotl limb blastema [39]. This data set consisted of samples taken at 0 hours, 3 hours, 6 hours, 12 hours, 1 day, 3 days, 5 days, 7 days, 10 days, 14 days, 21 days, and 28 days after the start of regeneration and had a total of 345,702,776 strand non-specific, single-end reads. Because of the large size of this data set and our goal of testing many different assemblers and parameter settings, we first restricted our analysis to data from three of the time points (6 hours, 14 days, and 28 days), which made up a total of 55,559,405 reads. We ran Trinity, Oases and SOAPdenovo-Trans on these data to produce over 100 different assemblies, each of which we scored using RSEM-EVAL. Trans-ABySS was not included due to some difficulties in running it.

Since we did not have a known axolotl transcript set, we were unable to use the reference-based measures we have discussed thus far to assess the RSEM-EVAL score’s effectiveness on these data. Therefore, to obtain an orthogonal measure of accuracy with which to validate the RSEM-EVAL score on this data set, we instead used alignments of the assembly contigs to the known protein sequences of the frog species *Xenopus tropicalis*. Specifically, we aligned the assemblies against the frog protein sequences with BLASTX [1] and calculated the the number of frog proteins that were recovered to various percentages of length by an axolotl contig (Additional file 1, Section 9). We found that, in general, the assemblies with higher RSEM-EVAL scores were those that were also considered better by comparison with the *Xenopus* protein set (Figure 9). Thus, the RSEM-EVAL score appears to be selecting the highest quality axolotl assemblies.

**Figure 9:**
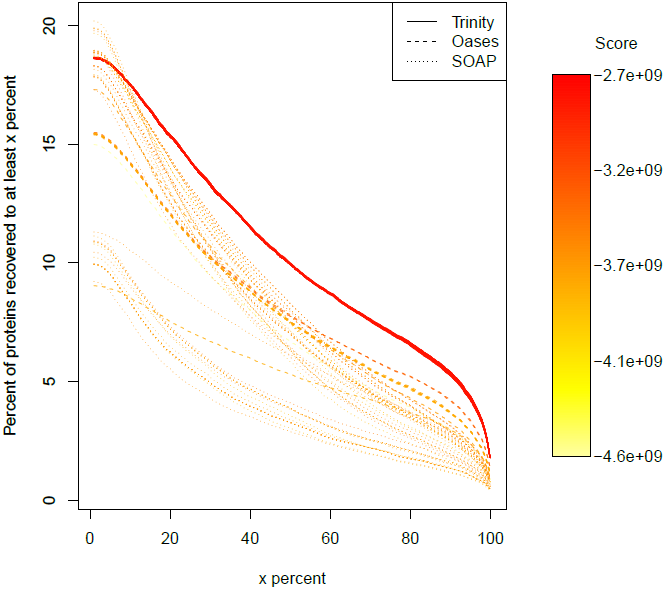
RSEM-EVAL scores and *Xenopus* protein recovery for the axolotl blastema transcriptome assemblies. The y-axis represents the percent of proteins with at least x percent of their length (x-axis) recovered by an axolotl contig. The curve for each assembly is colored according to its RSEM-EVAL score, with red representing the highest RSEM-EVAL score. The assembly with the curve closest to the upper-right corner is the best in terms of its comparison with the *Xenopus* protein set.

We then pooled all timepoints of the timecourse and built an assembly using the assembler (Trinity) and parameter set (`--glue_factor 0.01 --min_iso_ratio 0.1`) that maximized the RSEM-EVAL score on the subset described above. This assembly is publicly available at the DETONATE website. We compared this assembly to a published axolotl assembly [39]. We find that the new assembly is longer overall and has a larger N50 score than the published assembly (Additional file 1, Table S3).

As length-based measures may not be indicative of a higher quality assembly, we also evaluated the assemblies based on the number of expressed genes and the number of up-regulated differentially expressed genes (DE UP) at each timepoint in the the axolotl RNA-seq timecourse. To enable direct comparisons with the published assembly, we used data and methods identical to those in [39], which used a comparative technique that analyzes the axolotl transcripts in terms of their orthologous human genes. With the new RSEM-EVAL guided assembly, we identify both more expressed genes at each timepoint and more DE UP genes at each timepoint (Additional file 1, Figure S15). The majority of DE UP genes found in the published assembly are captured in the new assembly (608 of 888 = 68%), while only 39% (608 of 1576) of the DE UP genes found in the new assembly are captured in the published assembly. The new assembly identifies many new DE UP genes (968) not captured in the old published assembly.

Because transcription factors (TFs) are important for setting and establishing cell state [41], we further evaluated the list of TFs found in the new assembly that are not found in the published assembly across the axolotl RNA-seq timecourse. Prior results indicate that oncogenes are upregulated early in the timecourse [39]. Using the new assembly we identify two additional DE UP oncogenes (*FOSL1* and *JUNB*) that are not identified using the published assembly [39]. The prior assembly identified many genes associated with limb development and limb regeneration as being DE UP during the middle phase (3d-14d) of the timecourse [39]. The new assembly identifies additional limb development and limb regeneration genes during this middle phase such as *HOXA11*, *HOXA13*, *MSX1*, *MSX2*, and *SHOX*. *HOXA11* and *HOXA13* are important specifiers or markers of positional information along the proximal/distal and anterior/posterior axes of the limb [46]. *MSX1* and *MSX2* have been shown to be expressed in the axolotl blastema [17]. *SHOX* mutants in human exhibit short limbs and overall stature [7]. The identification of many more expressed and DE UP genes, a number of which have previous support for involvement in limb regeneration, suggests that the new assembly gives a more comprehensive view of the genes expressed in the axolotl.

## Discussion

### Related work

RSEM-EVAL is related to several recently developed methods for *de novo* genome and metagenome assembly. Although the evaluation of *de novo* genome assemblies is a relatively mature area of research [8, 9, 13, 23, 30, 32, 36, 43, 47], only recently has the need for statistically-principled methods been recognized [14]. Notably, Rahman and Pachter [33] developed CGAL, a principled probabilistic model–based method for evaluating genome assemblies without the ground truth. RSEM-EVAL and CGAL are closely related in that they both make use of the likelihood to score an assembly. However, RSEM-EVAL is necessarily more complex due to important differences between the tasks of transcriptome and genome assembly. In particular, with genome assembly it is generally assumed that all chromosomes are sequenced to the same sequencing depth, whereas with transcriptome assembly one has to consider variable sequencing depth because of widely-varying transcript abundances. Therefore, as we demonstrated through our experiments, use of the likelihood alone is suboptimal for evaluating transcriptome assemblies. In fact, even for *de novo* genome assembly evaluation, one can show that use of the likelihood alone is not optimal. For example, the likelihood criterion cannot distinguish between the “true” assembly and assemblies constructed by duplicating each contig of the “true” assembly *c* times. RSEM-EVAL corrects for this limitation by prior modeling of assemblies.

Because the task of *de novo* metagenome assembly is roughly equivalent to that of *de novo* transcriptome assembly, RSEM-EVAL also has close connections to methods developed for metagenomics. Metagenomic data are similar to transcriptomic data in that each distinct element of the population is present at varying multiplicities. Laserson et al. [20] showed that metagenome assembly can be improved by finding an assembly that maximizes a probabilistic model–based objective function. However, unlike RSEM-EVAL, their objective function does not handle multi-mapping reads appropriately, which are of great significance in *de novo* transcriptome assembly primarily because of alternative splicing. Clark et al. [5] developed ALE, a probabilistic model–based evaluator for both genome and metagenome assemblies that models the prior of each assembled contig by its 4-mer frequencies. ALE’s prior can be used to detect chimeric contigs formed by wrongly fusing two or more genomes, but it does not model an assembly’s complexity. In addition, ALE’s model does not take strain frequencies into consideration, which reduces its utility for evaluating transcriptome assemblies or metagenome assemblies in which the strains are present at highly variable frequencies.

### Limitations and future work

We stress that the correlation experiments presented in this study were primarily designed to evaluate the RSEM-EVAL score, not to determine which assembler is most accurate, in general. The versions of the assemblers used were not the most recent ones at the time of this writing and the parameter variations used were not necessarily those recommended by the assemblers. Nevertheless, with our selected assembler versions and parameter variations (which included the default settings for each assembler), our results provide some evidence that Trinity is more accurate than the other assemblers. To confirm this, future benchmarking will require use of the latest versions of these actively-developed assemblers, additional data sets, and more carefully selected parameters.

We also note a couple of important limitations regarding the use of RSEM-EVAL. First, it is critical that RSEM-EVAL be run on the same RNA-Seq data set that was provided to the assemblers as it assumes that an assembly is compatible with the data. Second, RSEM-EVAL should not be used if genome information is used during transcriptome assembly because its model is purely based on the RNA-Seq data and therefore might be misleading for genome-guided assemblies. Lastly, RSEMEVAL currently only focuses on contig assemblies constructed from single-end Illumina RNA-Seq data. We choose to focus on the Illumina platform as it is currently the most popular RNA-Seq platform and to restrict our methods to single-end reads and contig assemblies for simplicity. However, the methods we have presented can, in principle, be extended to paired-end data, scaffold assemblies, and other sequencing platforms, and we plan to do so in the near future.

Since RSEM-EVAL does not currently handle paired-end reads explicitly, the performance of assemblers may be different than that suggested by RSEM-EVAL because assemblers differ in their utilization of paired-end information. However, RSEM-EVAL can be used to evaluate the contigs of assemblies built with paired-end data with the workaround of providing the union of the two mates as one single-end data set to RSEM-EVAL. Although this is not ideal, in part because the two mates of a paired-end reads are not independent, we expect it will give reasonable results until RSEM-EVAL fully supports paired-end data and scaffolding.

In addition to handling paired-end data, we plan to extend our framework to support other sequencing platforms (e.g., Roche 454 sequencing, Ion Torrent and Pacific Biosciences), indel alignments, and correction for sequencing biases. We also plan to investigate remedies to RSEM-EVAL’s weakness of scoring assemblies with incorrectly fused contigs above the ground truth. Although it is unlikely that any reference-free measure can score the ground truth above all other assemblies, our random perturbation experiments suggest that RSEM-EVAL is permissive of assemblies that concatenate low-coverage contigs, perhaps because of its modeling of the “true” assembly with minimum overlap length 0.

## Conclusions

We presented DETONATE, a methodology and corresponding software package for evaluating *de novo* transcriptome assemblies that has the ability to compute both reference-free and reference-based measures. RSEM-EVAL, our reference-free measure, uses a novel probabilistic model–based method to compute the joint probability of both an assembly and the RNA-Seq data as an evaluation score. Since it only relies on the RNA-Seq data, it can be used to select a best assembler, tune the parameters of a single assembler, and guide new assembler design, even when the ground truth is not available. REF-EVAL, our toolkit for reference-based measures, allows for a more refined evaluation compared to existing reference-based measures. The measures it provides include recall, precision and *F*_1_ scores at the nucleotide and contig levels, as well as a *k*-mer compression score.

Experimental results based on both simulated and real data sets show that our RSEM-EVAL score accurately reflects assembly quality. Results from perturbation experiments that explored the local assembly space around the ground truth suggest that RSEM-EVAL ranks the ground truth among the locally highest scoring assemblies. In contrast, a score based only on the likelihood fails to rank the ground truth among its best scores, which highlights the importance of including a prior on assemblies as part of an evaluation score. Through correlation experiments we measured the similarity of the RSEM-EVAL scores to different reference-based measures. We find that, in general, the RSEM-EVAL score correlates well with reference-based measures. In contrast, the commonly used N50 measure does not correlate well, which suggests that it is inappropriate for evaluating transcriptome assemblies.

To demonstrate the usage of RSEM-EVAL, we assembled a set of contigs for the regenerating axolotl limb with its guidance. Evaluation of this new assembly suggests that it gives a more comprehensive picture of genes expressed and genes up-regulated during axolotl limb regeneration. Thus, RSEMEVAL is likely to be a powerful tool in the building of assemblies for a variety of organisms where the genome has not yet been sequenced and/or the transcriptome has not yet been annotated.

## Methods

### The “true” assembly according to DETONATE

As discussed in the Results above, both RSEM-EVAL and REF-EVAL rely on the concept of the “true” assembly of a set of RNA-Seq reads, which is the assembly one would construct if given knowledge of the true origin of each read. For each read, *r*, let *transcript*(*r*), *left*(*r*), and *right*(*r*), denote the transcript from which *r* truly originates, the leftmost (5’) position of *r* along the transcript, and the rightmost (3’) position of *r* along the transcript, respectively. We parameterize this notion of a “true” assembly by a length, *w*, which is the minimum overlap between two reads required for the extension of a contig. Given *w*, we define the sequence of a segment, [*start, end*], of a transcript, *t*, to be a “true” contig if there exists an ordered set of reads, (*r*_1_*, r*_2_*,…, r_n_*), such that, (i) *transcript*(*r_i_*) = *t, ∀i*, (ii) *right*(*r_i_*) – *left*(*r_i_*_+1_) + 1 *≥ w, ∀i < n*, (iii) *left*(*r*_1_) = *start*, (iv) *right*(*r_n_*) = *end*, and (v) [*start, end*] is maximal. The first four conditions ensure that the segment is completely covered by reads overlapping by at least *w* bases and the last condition ensures that one contig cannot be contained within another. The “true” assembly is then defined as the set of all “true” contigs. Such an assembly is the best theoretically achievable by a *de novo* assembler that requires at least *w* bases of overlap to merge two reads. The “true” assembly at minimum overlap length *w* = 0 is the best theoretically achievable assembly in that it represents all contiguously-covered segments of the transcript sequences.

### Overview of RSEM-EVAL

RSEM-EVAL models an RNA-Seq data set, *D*, consisting of *N* single-end reads, each of length *L*, and an assembly, *A*, consisting of *M* contigs, with the length of contig *i* denoted by *l_i_* and its sequence by *a_i_*. RSEM-EVAL also models the expected read coverage Λ = {λ*_i_*} of each contig’s parent transcript, where the expected read coverage of a transcript is defined as the expected number of reads that start from each valid position of the transcript, given the sequencing throughput, *N*. A transcript’s expected read coverage is proportional to its relative expression level (Additional file 1, Section 1).

A natural way to decompose the joint distribution of an assembly and the reads used to construct it, *P* (*A, D*), would be to (1) specify a prior distribution over the unobserved transcript sequences and their abundances, (2) model the generation of reads from these transcripts, and (3) model *A* as being the “true” assembly at minimum overlap length 0 of the reads in *D* (Additional file 1, Figure S1(a)). Unfortunately, this decomposition requires us to integrate out the unobserved transcript sequences in order to obtain the distribution *P* (*A, D*), and doing so is computationally infeasible.

Instead, RSEM-EVAL decomposes the joint distribution in an equally valid but more computationally convenient manner, as follows. (1) We specify a *prior* distribution, *P* (*A|*Λ), over the assembly *A*, given the expected read coverage of each contig’s parent transcript, Λ. (2) We model the generation of a set of reads *D* consistent with the assembly *A* and the expected read coverage Λ; this model defines the *likelihood P* (*D|A,* Λ), the probability of the reads *D* given the assembly A and the expected read coverage Λ. (3) Instead of specifying a concrete distribution over the number of contigs *M* and the expected read coverage Λ, we approximately integrate out these variables using the *Bayesian information criterion* (BIC) [38].

Based on this decomposition, the RSEM-EVAL score, log *P* (*A, D*), can be expressed as a sum of three terms, the assembly prior, the likelihood, and a BIC term (Equation 1 in the Results). These three terms are detailed in the next three subsections. The relationship between the “natural” decomposition and the RSEM-EVAL model is further discussed in Additional file 1, Section 2.

### RSEM-EVAL’s assembly prior component

RSEM-EVAL’s prior distribution over assemblies is based on a simple parametric model of the transcriptome and the reads, together with the concept of a “true” assembly as we have described. Details are as follows.

First, a key assumption of the prior model is that each contig is generated independently. This assumption is useful, even though it is not satisfied in practice since multiple contigs can be generated from the same transcript. With this assumption, we may express the the prior as

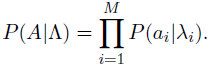

Second, each contig’s parent transcript is modeled as follows. (1) The transcript’s length follows a negative binomial distribution, the parameters of which may be estimated from a known transcriptome. (2) The transcript’s sequence follows a uniform distribution, given the transcript’s length. (3) For each position in the transcript, the number of reads starting at that position follows a Poisson distribution (mean = expected read coverage), independently, given the transcript length and the expected read coverage.

Third, each contig’s distribution is induced from the distribution of its parent transcript, as follows. Imagine that we repeat the following steps until we have a large enough number of contigs in the bag. (1) A transcript and its reads are generated as above. (2) The “true” assembly at minimum overlap length *w* = 0 is constructed from these reads. (3) All contigs in the “true” assembly are put into the bag. The frequency of contigs in the resulting bag defines our per-contig prior *P* (*a_i_|λ_i_*).

The above specification leads to the following functional form for the prior:

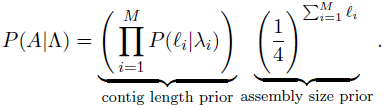

One can work out a dynamic programming algorithm to compute *P* (ℓ*_i_|λ_i_*), by means of which the prior can be computed efficiently (Additional file 1, Section 3).

The practical contribution of the prior is as follows. Each term of the contig length prior, *P* (ℓ*_i_|λ_i_*), penalizes contigs with aberrant lengths that are not likely given the expected read coverage of their parent transcripts. The assembly size prior, 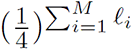, imposes a parsimony preference on the total length of an assembly and pushes the RSEM-EVAL score to favor assemblies using fewer bases to explain the RNA-Seq data.

### RSEM-EVAL’s likelihood component

For modeling the RNA-Seq reads, we build off of the model used by RSEM [21, 22] for the task of transcript quantification. The RSEM model provides a probability distribution for an RNA-Seq data set, *D*, given known transcript sequences, *T*, and relative abundances of those transcripts encoded by the parameters, Θ. Given this model, RSEM uses the Expectation-Maximization (EM) algorithm for efficient computation of the ML estimates for Θ. Unfortunately, we cannot directly use the likelihood under the RSEM model because (1) we do not observe the full-length transcript sequences and (2) we require that the RNA-Seq reads are consistent with an assembly in that they completely cover the contigs. Nevertheless, the RSEM model can be used as part of a generative process that results in a proper probability distribution over *D*, given an assembly, *A*. This process follows a simple two-step rejection procedure. (Step 1) Generate a set of reads, *D*′, according to the RSEM model with the contigs in *A* treated as full-length transcripts. (Step 2) If the reads in *D*′ completely cover the contigs, then set *D* = *D*′, otherwise go back to Step 1. This process results in the following form for RSEM-EVAL’s likelihood:

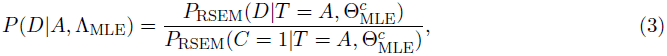

where *P*_RSEM_ denotes a probability under the RSEM model and *C* = 1 denotes the event that every position in the assembly is covered by reads that overlap with each other by at least *w* bases. 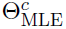 denotes the equivalent parameter values of the RSEM model given the ML expected read coverage values, Λ_MLE_. We refer to the denominator of Equation 3 as the *likelihood correction term*. This term can be calculated efficiently (Additional file 1, Section 4).

### RSEM-EVAL’s BIC penalty component

The BIC penalty is proportional to the product of the number of free parameters and the logarithm of the size of the data. The free parameters consist of the expected coverage of each contig, plus one extra parameter for the expected number of reads from RSEM’s model, for *M* + 1 parameters in total. The data size is *N*, which represents the number of reads. The BIC penalty imposes a parsimony preference on the total number of contigs in an assembly.

### The inference algorithm used to compute the RSEM-EVAL score

We use the following heuristic inference algorithm to calculate the RSEM-EVAL score:

1. Learn Θ_MLE_ using RSEM, treating the input assembly *A* as the true transcript set.
2. Convert Θ_MLE_ into Λ_MLE_ via the formula 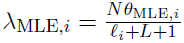, for *i >* 0.
3. Calculate the RSEM-EVAL score, log *P* (*A, D*), using Λ_MLE_ and Equation 1.

In step 1, RSEM requires use of a short-read alignment program to generate candidate alignments of each read to the assembly [21, 22]. The Bowtie aligner (v0.12.9) [19] was used for all experiments except the random perturbation experiments. Bowtie was called through RSEM with RSEM’s default Bowtie parameters. For the random perturbation experiments, since the perturbations were subtle, we chose to use a more sensitive aligner, GEM (binary pre-release 3) [25]. In order to produce alignments with similar criteria as Bowtie, we first extracted the first 25bp (seed) of every read and aligned the seeds using GEM with `-q ignore --mismatch-alphabet ACGNT -m 2 -e 0 --max-big-indel-length 0 -s 2 -d 200 -D 0` options set for `gem-mapper`. Lastly, we filtered any alignments for which the seed could not be extended to the full read length.

In step 2, we do not use the contig level read coverage, 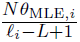 to approximate the expected read coverage of its parent transcript, *λ_i_*, because it is likely to overestimate *λ_i_*, especially for short contigs. Instead, to partially correct for this overestimation, we adjust the denominator in this estimator to *l_i_* + *L* + 1 to account for the fact that no reads started within *L* bases of the start positions of the reads making up the contig (otherwise, the contig would have been extended, since our model considers the assembly to have been created using minimum overlap length *w* = 0).

### REF-EVAL’s estimate of the “true” assembly

REF-EVAL estimates a set of “true” contig sequences from a given set of transcripts and RNA-Seq reads using the following procedure:

1. Align the reads against the transcript sequences using RSEM.
2. For each alignable read, sample one of its alignments based on the posterior probability that it is the true alignment, as estimated by RSEM. The set of alignments for each read includes the null “alignment” that the read comes from background noise.
3. Treat the sampled alignments as true alignments and compute the “true” contigs with minimum overlap length *w* = 0.

### REF-EVAL’s contig- and nucleotide-level measures

Given a set of ground truth reference sequences, REF-EVAL provides assembly recall, precision, and *F*_1_ scores at two different granularities. Recall is the fraction of reference elements (contigs or nucleotides) that are correctly recovered by an assembly. Precision is the fraction of assembly elements that correctly recover a reference element. The *F*_1_ score is the harmonic mean of recall and precision:

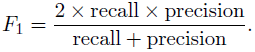

Because the *F*_1_ score is a combination of both recall and precision, it gives a more balanced view of an assembly’s accuracy than either precision or recall does alone.

REF-EVAL provides these measures at two different granularities: contig and nucleotide. Taking recall as an example, the measures at the two levels can be summarized as follows. For both levels, we first compute all significant local alignments between the assembly contigs and the reference sequences, using BLAT [16]. The contig level measurement [29, 34, 35] counts the number of correctly recovered reference sequences after a one-to-one mapping is established between contigs and reference sequences. At the nucleotide level, the recall measure instead counts the number of correctly recovered nucleotides based on the alignments between the assembly and the ground truth sequences [2, 37]. The precision measures for both levels are calculated similarly by switching the roles of the assembly contigs and reference sequences.

In detail, REF-EVAL defines the contig recall as follows. For a reference sequence to be correctly recovered, at least 99% of its sequence must be identical to that of the contig to which it is aligned, and the total number of insertions and deletions in the alignment between the two must be no more than 1% of the length of either sequence. Each percentage is computed relative to the length of the contig or the length of the reference sequence, whichever is most stringent. Under these criteria, multiple reference sequences can be recovered by the same contig. To handle this issue, we define a bipartite graph in which the vertices are the assembly contigs and the reference sequences, and the edges correspond to alignments that meet the above criteria. The contig recall is the cardinality of the maximum cardinality matching of this graph.

In detail, REF-EVAL defines the nucleotide recall as follows. A nucleotide in a reference sequence is considered to be correctly recovered if the corresponding nucleotide in the contig alignment selected for that position is identical. To handle the issue of multiple local alignments overlapping a given reference position, we select a single alignment for each position by picking alignments in order of their marginal contribution to the nucleotide recall, given the alignments that have already been selected.

Algorithms to compute the contig and nucleotide measures are given in Additional file 1, Sections 6 and 7.

### REF-EVAL’s ***k***-mer compression score

The *k*-mer compression score (KC score) is a combination of two measures, weighted *k*-mer recall (WKR) and inverse compression rate (ICR) (Equation 2). Details about the WKR and ICR measures are provided below.

The WKR measures an assembly’s recall of the *k*-mers present in the reference sequences, with each *k*-mer weighted by its relative frequency within the reference transcriptome. It has several advantages over the contig and nucleotide level recall measures. First, unlike the nucleotide measure, it takes into account connectivity between nucleotides, but is not as stringent as the contig measure because it only considers connectivity of nucleotides up to *k* - 1 positions apart. Second, it is biased towards an assembler’s ability to reconstruct transcripts with higher abundance, which are arguably more informative for evaluation because there is generally sufficient data for their assembly. Lastly, it is relatively easy to compute because it does not require an alignment between the assembly and the reference sequences.

To compute the WKR, the relative abundances of the reference elements are required. These abundances may be estimated from the RNA-Seq data used for the assembly, and REF-EVAL uses RSEM for this purpose. Given the reference sequences and their abundances, a *k*-mer occurrence frequency profile, *p*, is computed, with individual *k*-mer occurrences weighted by their parent sequences’ abundances: for each *k*-mer *r*, we define

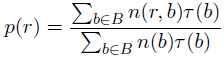

where *B* is the set of reference sequences, and for each reference sequence *b* in *B*, *n*(*r, b*) is the number of times the *k*-mer *r* occurs in *b*, *n*(*b*) is the total number of *k*-mers in *b*, and *τ* (*b*) is the relative abundance of *b*. Letting *r*(*A*) be the set of all *k*-mers in the assembly, WKR is defined as

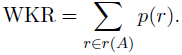

Since recall measures only tell half of the story regarding accuracy, the KC score includes a second term, the ICR, which serves to penalize large assemblies. We define the ICR of an assembly as

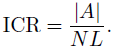

As this term’s name suggests, we can view an assembler as a “compressor” for RNA-Seq data and the assembly it outputs as the “compressed” result. The uncompressed data size is the total number of nucleotides in the data, *N L*, and the compressed result size is the total length of the assembly, *|A|*. The smaller the ICR value, the more efficient the assembler is in compressing the data. We chose to use ICR over other possible precision-correcting schemes (e.g., *F*_1_ score) because of important mathematical connections between the RSEM-EVAL and KC scores (Additional file 1, Section 8). Thus, as we showed with our experiments, the reference-based KC score provides some additional intuition for what the reference-free RSEM-EVAL score is measuring. The theoretical links between the two scores suggest that the KC score would benefit from some additional terms (Additional file 1, Section 8), however, in keeping with our goal of simplicity, we restricted the KC score to the two most dominant terms.

### RNA-Seq data used in the perturbation experiments

Both the random and the guided perturbation experiments (see Results) use a simulated set of RNA-Seq reads. We simulated these reads from a mouse transcript set (Ensembl Release 63 [11]), using RSEM’s [21,22] RNA-Seq simulator and simulation parameters learned from the first mates of a set of real paired-end mouse data (SRA accession SRX017794 [42]). The resulting simulated data set contained around 42 million strand non-specific, 76bp single-end reads. For reasons of computational efficiency, in the random perturbation experiments, we only used reads from transcripts on chromosome 1, and we constructed the ground truth accordingly. This resulted in 1,843,797 reads that formed 10,974 contigs in the ground truth assembly.

For our guided perturbation experiments, we additionally used a real data set consisting of the first mates of the mouse data previously mentioned. This real data set also contained around 42 million strand non-specific, 76bp single-end reads. The true origin of each read in the real data set was estimated using REF-EVAL.

### Construction of randomly-perturbed assemblies

The random perturbation experiments (see Results) compare the “true” assembly’s RSEM-EVAL score to the scores of numerous perturbed variants of this assembly. These perturbed assemblies were constructed as follows.

Substitution assemblies (randomly-perturbed assemblies with substitution mutations) were generated by randomly and independently substituting each base of the ground truth at a specific substitution rate. Fusion assemblies were generated by repeatedly joining two randomly selected contigs until the specified number of fusion events was reached. For each join, if the two selected contigs shared a common sequence at their ends to be joined, the contigs were overlapped such that the shared sequence only appeared once in the fused contig. To generate a fission assembly, each contig position at which two reads were merged in the ground truth was independently selected as a fission point at the specified fission rate. If reads overlapped at a fission point, the overlapped segment was made to appear in both contigs after the fission. To generate an indel-perturbed assembly, insertions and deletions were introduced across the ground truth assembly according to the specified indel rate. The length of each indel was sampled from a geometric distribution with mean 3, to model the short indels occasionally observed in high-throughput sequencing data.

For substitution and indel assemblies, the mutation strength was the mutation rate per contig position, which was varied among 1e-6, 1e-5, 1e-4, 1e-3 and 1e-2. For example, in the substitution experiment, at mutation rate 1e-3, a substitution was introduced at each position of each contig with probability 1e-3. The fission experiments used the same range of mutation rates, but with the rate defined as per pair of immediately adjacent and overlapping reads. For fusions, the mutation strength was the number of fusion events, which ranged among 1, 10, 100, 1000, 10000.

### RNA-Seq data used in the correlation experiments

For our experiments measuring the correlation between the RSEM-EVAL score and reference-based measures we used four data sets: the simulated and real mouse, strand non-specific, single-end RNA-Seq data sets used in the perturbation experiments, and the strand-specific mouse and yeast data sets from Grabherr et al. [12]. The entirety of each data set was used, except for the simulated data set, from which we used roughly half of the simulated reads (reads from transcripts in chromosome 1 to 10) because of computational efficiency reasons. For each assembler, we used a variety of parameter settings (Additional file 2), resulting in a total of 211 assemblies for each of the strand non-specific data sets and 159 assemblies for each of the strand-specific data sets (Table 1). Because the versions of SOAPdenovo-Trans and Trans-ABySS we used did not take into account strand-specificity, they were not run on the two strand-specific data sets. The reference-based measures were computed relative to the estimated “true” assemblies at minimum overlap length *w* = 0. These assemblies were estimated using the mouse Ensembl annotation (release 63) for the three mouse data sets and the PomBase [44] *S. pombe* annotation (v09052011) for the yeast data set.

**Table 1.**
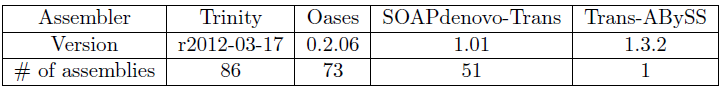
The assemblers and the number of assemblies generated by them (using different parameter settings) for the correlation experiments. SOAPdenovo-Trans and Trans-ABySS were not run on the strand-specific data sets.

| Assembler | Trinity | Oases | SOAPdenovo-Trans | Trans-ABYSS |
| --- | --- | --- | --- | --- |
| Version | r2012-03-17 | 0.2.06 | 1.01 | 1.3.2 |
| # of assemblies | 86 | 73 | 51 | 1 |

### Software availability

The DETONATE software is freely available from its website: http://deweylab.biostat.wisc.edu/detonate.

## List of abbreviations

BIC: Bayesian information criterion; ML: maximum likelihood; KC score: *k*-mer compression score; WKR: weighted *k*-mer recall; ICR: inverse compression rate; EM: Expectation-Maximization

## Author’s contributions

CND conceived the research. BL, NF, and CND designed RSEM-EVAL and REF-EVAL. BL implemented RSEM-EVAL. NF implemented REF-EVAL and assembled the DETONATE package. BL and NF performed the analysis. YB, MC, JAT, and RS contributed the axolotl data and performed axolotl-related analysis. BL, NF, RS, and CND wrote the manuscript. All authors read and approved the final manuscript.

## Acknowledgements

We thank Lior Pachter, Sreeram Kannan, and Marcel H. Schulz for helpful discussions. This work was supported by the National Institutes of Health (grants R01HG005232 and T15LM007359).

